# scRNA-seq Reveals A Macrophage Subset That Provides A Splenic Replication Niche For Intracellular *Salmonella*

**DOI:** 10.1101/2021.01.04.425175

**Authors:** Dotan Hoffman, Yaara Tevet, Gili Rosenberg, Leia Vainman, Aryeh Solomon, Shelly Hen-Avivi, Noa Bossel Ben-Moshe, Roi Avraham

## Abstract

Interactions between intracellular bacteria and mononuclear phagocytes give rise to diverse cellular phenotypes that may determine the outcome of infection. Recent advances in single cell RNA-seq (scRNA-seq) have identified multiple subsets within the mononuclear population defined by unique molecular features, but the implications to their function during infection is unknown. Here, we applied high resolution kinetic analysis using microscopy, flow cytometry and scRNA-seq to survey the mononuclear niche of intracellular *Salmonella* Typhimurium (*S*.Tm) during early systemic infection in mice. We describe an eclipse like growth kinetics in the spleen, with a first phase of bacterial control mediated by tissue resident red pulp macrophages. A second phase involved bacterial growth mediated by intracellular replication within a macrophage population we termed CD9 macrophages, that originate from non-classical monocytes. *Nr4a1^e2−/−^* mice, specifically depleted of non-classical monocytes but not other mononuclear cells, are more resistant to *S*.Tm infection. Our study underscores a cell-type specific host-pathogen interaction that determines early infection growth dynamics and has implications to the infection outcome of the entire organism.

## Introduction

Encounters between pathogenic bacteria and mononuclear phagocytes during early systemic infection can determine the capacity of the host to eliminate the invading pathogen or the ability of bacteria to establish a successful infection (Mills and Avraham, 2017). One of the major sites for clearance of infection (Carter and Collins, 1974), the spleen is host to multiple mononuclear phagocyte populations: 1) Tissue resident macrophages; red pulp, metallophilic and marginal zone macrophages, are sessile and perform tissue-specific functions (De Jesus et al., 2008; Kirby et al., 2009). 2) Circulating classical monocytes from the blood constantly replenish the monocyte pool in the spleen and also convert into non-classical (NC) monocytes (Yona et al., 2013). 3) A reservoir of undifferentiated splenic monocytes that rapidly respond to inflammatory events (Swirski et al., 2009). Early after infection, tissue-resident macrophages respond and engage invading pathogens followed by recruitment of classical monocytes from the blood that differentiate into effector cells and allow efficient clearance of the pathogen (Serbina et al., 2008). Within the tissue, these macrophages and monocytes are heterogeneous population; a taxonomy of discrete cell types with continuous transitions of cell differentiation and activation states (Ginhoux and Guilliams, 2016; Trzebanski and Jung, 2020). The heterogeneous nature of mononuclear phagocyte populations suggests that their interaction with intracellular bacteria is likely to result in a variety of different, complex phenotypes.

Studies using flow cytometry and microscopy indicate that intracellular infection outcome is dependent on the identity of the infected monocytes/macrophage and the environmental context within the tissue (Italiani and Boraschi, 2014). In the lung, *Mycobacterium tuberculosis* is controlled by interstitial macrophages while resident alveolar macrophages are permissive to intracellular replication (Huang et al., 2018). In the gut, *Salmonella enterica* serovar Typhimurium (*S*.Tm) can survive within CD18^+^ phagocytes, which allow it to traverse the epithelial barrier and disseminate to the spleen (Vazquez-Torres et al., 1999). During systemic infection, *S*.Tm are found in the spleen within tissue-resident macrophages; red pulp macrophages (Rp Mϕ) identified as F4/80^+^ and marginal zone macrophages identified as Siglec1^+^ cells (Geddes et al., 2007; Salcedo et al., 2001). *Streptococcus pneumonia* bacteria are found replicating within resident CD169^+^ metallophilic macrophages (Ercoli et al., 2018), while *Listeria monocytogenes* induce the early necroptotic death of resident Kupffer cells in the liver (Blériot et al., 2015). Furthermore, recruitment of Ly6C^high^ inflammatory monocytes to the intestine restricts *S*.Tm infection (Tam et al., 2014), and during *Listeria* infection Ly6C^+^ monocytes differentiate into iNOS-producing monocyte-derived cells (Menezes et al., 2016). Thus, the identity of monocyte/macrophages can provide the basis for our understanding of the molecular details and outcome of intracellular infection. However, classification of the cell populations using *a priori*-defined cell surface markers is often controversial, and the assignment of different cell types and their function remains challenging.

Recent advances in single-cell RNA-sequencing (scRNA-seq) allow breakdown of complex tissues into cell types which have revolutionized our ability to characterize the variety of cells within the immune compartment (Hashimshony et al., 2012; Jaitin et al., 2014). Applied to mononuclear phagocyte populations, scRNA-seq revealed rich biology of cellular subsets that far exceeds our knowledge of this cellular compartment (Guilliams et al., 2018). In the lung, two interstitial macrophage subsets were identified based on Lyve-1 and MHCII expression (Chakarov et al., 2019), and patrolling NR4A1-dependent monocytes were reported to replenish the pool of one interstitial macrophages subset with features of antigen-presenting cells (Schyns et al., 2019). In the skin, three tissue resident macrophage subsets were associated with regeneration and surveillance of local nerves (Kolter et al., 2019). In the heart, resident macrophages were classified based on Tim4 expression, with monocytes contributing to replenish this population after cardiac infarction (Dick et al., 2019). Within adipose tissue, Trem2 defines lipid-associated macrophages that are activated upon tissue lipid loss (Jaitin et al., 2019). In the central nerve system, a monocyte subset activated by lipopolysaccharides (LPS) or interferon-γ is classified by Cxcl10 and represent pathogenic effector cells (Giladi et al., 2020). Despite increasing knowledge of diverse mononuclear phagocyte populations, we currently lack an understanding how these translate to their function during intracellular infection. What is needed is a comprehensive analysis of the intracellular niche within the different monocytes and macrophages, that will provide a fundamental understanding of infection biology.

Here, we utilized scRNA-seq to test whether and how mononuclear phagocyte populations in the spleen change and respond during systemic infection with *S.*Tm. We aimed to decipher the mechanistic underpinnings of different cellular subsets that determine the phenotype of intracellular infection, and to link the diversity of mononuclear subsets to infection outcomes in vivo.

## Results

### Early systemic infection with *S*.Tm followed eclipse-like growth dynamics

To characterize the kinetics of early systemic infection, we inoculated 8 weeks old female C57BL/6J mice intravenously (i.v.) with *S*.Tm (1×10^7^ CFU). We collected spleens at different time points after infection and assessed bacterial load by plating the spleen homogenate on selective agar plates. Already one-hour post-infection (hpi), about 90% of the inoculum was found in the spleen (**Fig. 1A**). Subsequently, the bacterial load in the spleen followed eclipse-like infection dynamics (ELID) comprising two discrete time windows. In the first phase (<8 hpi), *S*.Tm infection was controlled and the bacterial load decreased. The second phase (>8 hpi) was characterized by rapid *S*.Tm replication. To investigate the mononuclear phagocyte diversity during ELID, we analyzed spleens from infected mice and stained with antibodies to exclude lymphocytes and neutrophils (Lin^−^), and gated using CD11b and F4/80 (**fig. S1A**). We detected three populations: CD11b^−^ F4/80^+^ (Rp Mϕ), CD11b^+^ F4/80^−^ (monocytes and dendritic cells), and CD11b^+^ F4/80^+^ (monocyte-derived macrophages) which appeared in infected mice only (**Fig. 1B**). We observed changes in the monocytes/macrophage populations that mirrored ELID: the first phase of ELID involved reduction in Rp Mϕ, and an increase in monocyte-derived macrophages in the second phase. To test whether these cells contain intracellular bacteria, we infected mice with an *S*.Tm strain that constitutively express an episomal fluorescent protein (*S*.Tm-GFP). Flow cytometry analysis showed that 24 hpi, intracellular *S*.Tm were found mostly within macrophages (F4/80^+^ GFP^+^) but not monocytes (CD11b^+^F4/80^−^GFP^+^) (**Fig. 1; C and D**), indicating that these cells are the main cellular niche for *S*.Tm during systemic infection, as was previously shown (Geddes et al., 2007; Salcedo et al., 2001).

**Figure 1:**
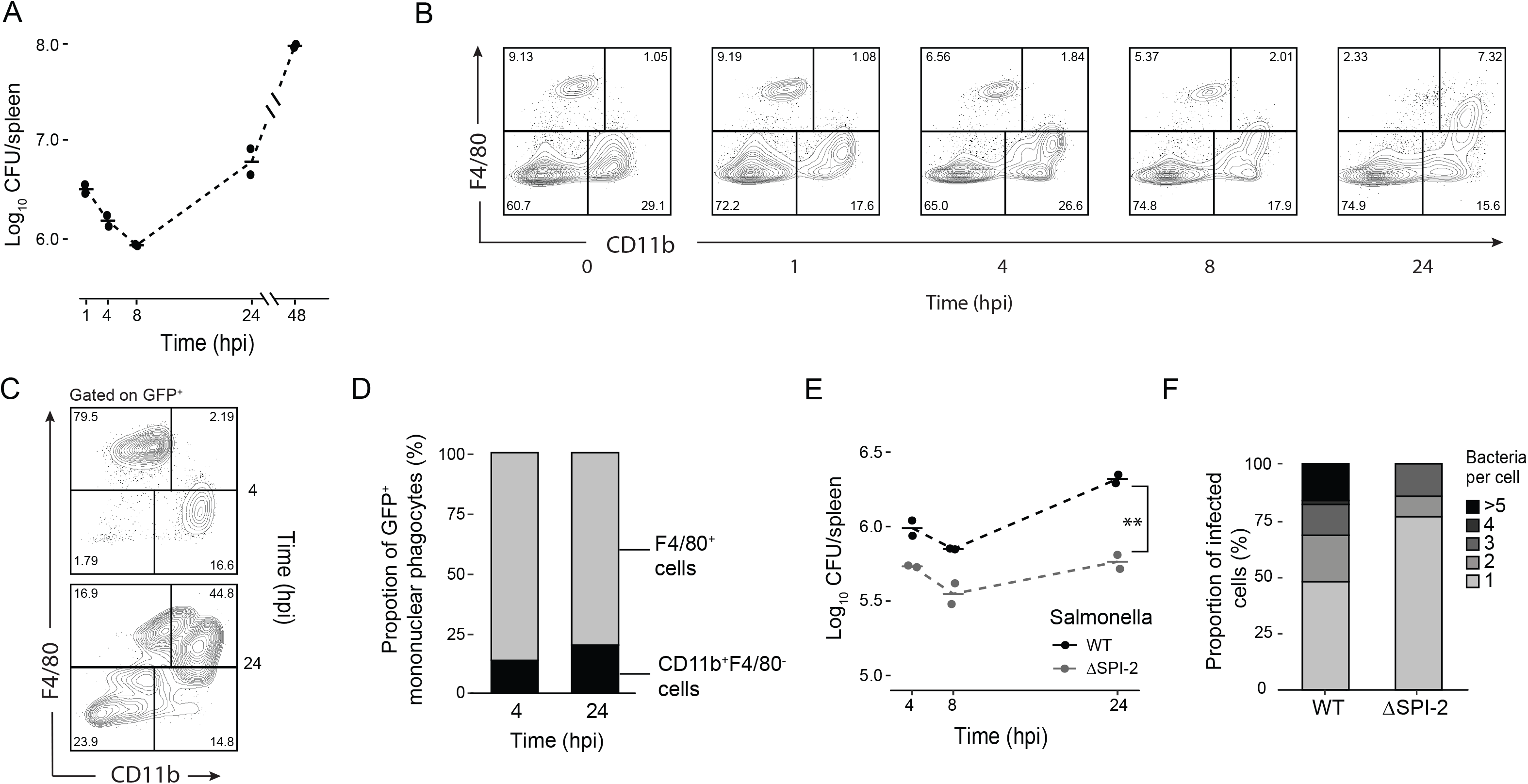
Early systemic infection with *S*.Tm followed ELID. (**A+B**) Mice were infected i.v. with *S*.Tm, and CFU was measured from spleens at 1, 4, 8, 24 and 48 hpi (**A**; n=2 per time point). Using flow cytometry, cells were gated to exclude Cd19, Cd3e, NK1.1 and Ly6g positive cells (Lin-) and analyzed for CD11b and F4/80 expression (**B**). (**C+D**) Mice were infected i.v. with *S*.Tm-GFP and spleens were harvested at 4 hpi (n=4) and 24 hpi (n=4). Infected cells were analyzed by flow cytometry using the GFP signal of the bacteria (**C**). Calculated ratios of GFP^+^ cells between F4/80^+^ and Cd11b^+^ F4/80^−^ populations are shown (**D**). (**E**) Mice were infected i.v. with either WT (black) or ΔSPI-2 (gray) *S*.Tm strains, and CFU was measured from spleens at 4, 8 and 24 hpi (n=2 per time point). (**F**) Mice were infected i.v. with either WT or ΔSPI-2 GFP-expressing strains. Twenty-four hpi, spleens were harvested, analyzed by flow cytometry and Cd11b^+^F4/80^+^ macrophages were sorted and individually plated on LB agar (n=94 in WT, n=34 in ΔSPI-2) to determine the number of CFU per cell. [(**A**) and (**E**)] Results are presented as median (line) and are representative of two independent experiments. (**E**) ** *P* < 0.01 using two-way ANOVA.

To assess whether ELID was a result of intracellular or extracellular replication of the pathogen within mononuclear phagocytes, we used an *S*.Tm mutant strain lacking the *ssaV* gene, encoding a structural protein of the needle complex used for type 3 secretion system located on the *Salmonella* pathogenicity island 2 (SPI-2). *S*.Tm mutants lacking *ssaV* are unable to secrete effectors through the SPI-2 apparatus (ΔSPI-2) (Hensel et al., 1998), which is essential for intracellular survival and replication of *S*.Tm within macrophages (Cirillo et al., 1998). We challenged mice with either wild type (WT) or a ΔSPI-2 *S*.Tm and assessed bacterial load in the spleen. Unlike WT, ΔSPI-2 mutant had significantly reduced recovery and replication in the spleen between 8-24 hpi (**Fig. 1E, fig. S1B**), suggesting that during the second phase, ELID requires intracellular replication. To further support this observation, we sorted single infected macrophages to enumerate single cell CFU (scCFU). We found that more than 50% of WT infected cells contained multiple replicating bacteria while the majority of ΔSPI-2 infected macrophages harbored a single intracellular bacterium (**Fig. 1F**). Thus, *S*.Tm growth follows ELID, which involves replication within macrophages mediated by SPI-2 and changes in the mononuclear phagocyte populations.

### scRNA-seq revealed three concomitant splenic macrophage populations during ELID of *S*.Tm

We next sought to decipher functional subsets of mononuclear phagocytes that are involved in ELID of *S*.Tm in the spleen. We infected mice with WT or ΔSPI-2 strains and gated for CD11b^+^F4/80^−^ and CD11b^−^F4/80^+^ populations in naïve mice, and CD11b^+^F4/80^+^ and CD11b^−^F4/80^+^ populations in mice infected with WT or ΔSPI2 (**Fig. 2A**). We did not gate on CD11b^+^F4/80^−^ in the infected mice as this population does not contain intracellular bacteria (**Fig. 1D**). Using fluorescently activated cell sorting (FACS), we sorted single cells from the gated populations and applied scRNA-seq (Jaitin et al., 2014) to characterize mononuclear phagocyte populations before and after infection. A total of 1982 cells from both uninfected (naïve) and infected mice passed quality control filters (**fig. S2A**). To define cell types according to the scRNA-seq data, we applied a MetaCell algorithm that groups cells with similar transcriptional profiles (Baran et al., 2019), producing a map of 27 MetaCells (MC) (**Fig. 2B**). To assign MCs to distinct cell types or activation states, we performed correlation analysis and identified MCs with similar transcriptional states (**fig. S2B**), which collapsed MCs to ten cell types with distinct gene expression program and marker genes (**Fig. 2C** and **fig. S2C)**. In naïve mice, MCs of the CD11b^+^ F4/80^−^ population were composed of dendritic cells (*MHCII*^+^*Cd74*^+^), classical monocytes (*Ly6c2*^+^*Ccr2*^+^), NC monocytes (*Nr4a1^+^Ace*^+^*Spn*^+^) that may derive from classical monocytes in the tissue (Olingy et al., 2017), and a small fraction of NK cells (*Nkg7*^+^*Gzma*^+^) (**fig. S2D**). MCs retrieved from the CD11b^−^F4/80^+^ population were annotated as Rp Mϕ in naïve mice (*Mertk*^+^*Mrc1*^+^) and in infected mice (*Marco*^+^*Cxcl9*^+^). Interestingly, in CD11b^+^ F4/80^+^ cells from the infected mice we identified two distinct MC populations. The first population, *Ly6c2*^+^*Nos2*^+^*Chi3l3*^+^ cells is reminiscent of iNOS Mϕ derived from Ly-6C^high^ classical monocytes (Menezes et al., 2016; Serbina et al., 2003). The second population identified by the expression of *Ly6c2*^−^ *Ly6i*^+^*Cd9*^+^, we termed CD9 Mϕ (**Fig. 2D)**. Noteworthy, infection with either WT or ΔSPI-2 *S*.Tm did not alter the MC identities following infection, but only their activation state (**fig. S2; E and F**), and were annotated as the same cell type for further analysis.

**Figure 2:**
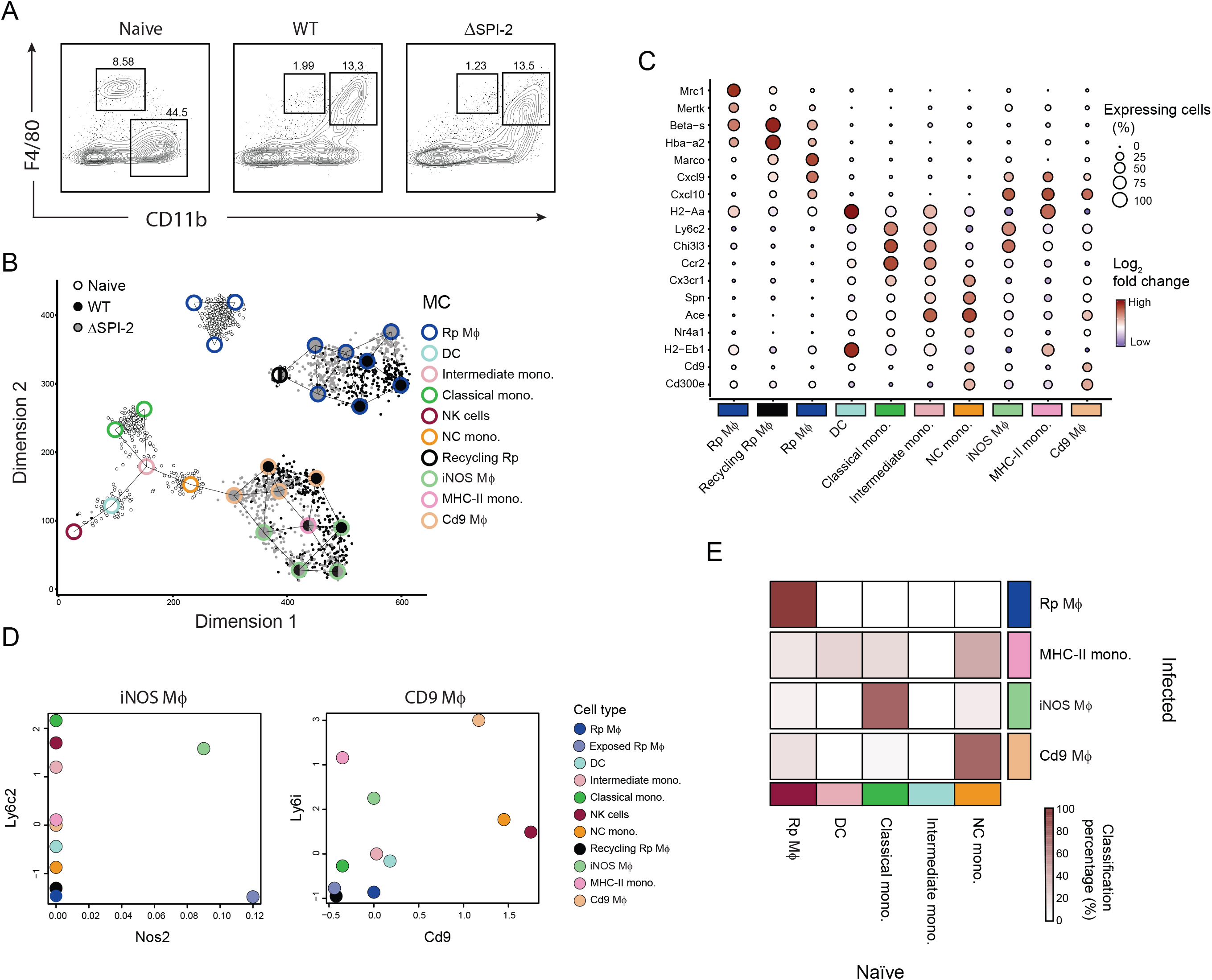
scRNA-seq revealed three concomitant splenic macrophage populations during ELID of *S*.Tm. (**A**) Mice were infected i.v. with *S*.Tm and 24 hpi spleens were harvested and analyzed by flow cytometry using CD11b and F4/80 antibodies. (**B**) Single cells from gated populations in A were sorted by FACS and analyzed by scRNA-seq. Shown is a forced layout map of single cells (dots) and MCs (large circles), with infection conditions indicated [naïve mice (white, n=631 cells), mice infected with WT (black, n=643 cells) or a ΔSPI-2 mutant (gray, n=658 cells)]. Cell type annotations are indicated by outline color of the MC, infection status of the MC by fill color and similarity between MCs are indicated by connecting nodes. (**C**) Marker gene annotations presented as the percentage of cells expressing the indicated genes in each MC (shown as the size of the circle) and the relative log2 fold change of the gene expression in each MC (shown as the color of the circle). (**D**) Expression analysis across all MCs of genes used as markers for iNOS Mϕ (*Ly6c*, *Nos2*) and CD9 Mϕ (*Cd9*, *Ly6i*). (**E**) *k*NN classification of cell types presented as percentage of infected cells classified to cell types in the naïve sample. Color bar indicates percentage of cells of each cell type in the infected mice classified to cell types from the naïve mice.

In order to align mononuclear populations before and after infection that can suggest a progeny-product relationship, we applied a *k*-nearest neighbor (*k*NN) classification algorithm to link single cells from the infected mice to the MCs in the naïve mice. As expected, Rp Mϕ following infection were matched mostly with Rp Mϕ MC in the naïve, and iNOS-Mϕ were aligned with classical monocytes in the naïve sample. Importantly, CD9 Mϕ in the infected sample were classified mostly to NC monocytes (**Fig. 2E**).

Thus, scRNA-seq analysis suggested that ELID of *S*.Tm is mirrored by changes in the macrophage landscape, underscored by the putative emergence of a CD9 Mϕ population from NC monocytes.

### Functionally distinct activation programs of iNOS Mϕ monocytes and CD9 Mϕ

We next followed the dynamics of macrophage populations after *S*.Tm infection using flow cytometry by staining for the cell surface molecules Ly6C and the tetraspanin receptor CD9 that are, according to our scRNA-seq analysis above, differentially expressed by iNOS Mϕ and CD9 Mϕ, respectively. This phenotypic characterization confirmed the existence of two distinct cell populations: iNOS Mϕ (Lin^−^CD11b^+^F4/80^+^CD64^+^Ly6C^+^CD9^−^) and CD9 Mϕ (Lin^−^CD11b^+^F4/80^+^CD64^+^Ly6C^−^CD9^+^) (**Fig. 3A and fig. S3A**). To further characterize these populations, we performed bulk RNA-seq on 1000 sorted cells from each population and curated a list of 310 differentially expressed genes (**Fig. 3B, fig. S3B and table S1**, *Q*<0.05). The expression signature of marker genes detected by the scRNA-seq matched the expression profiles of cells sorted using CD9 and Ly6C markers (genes indicated to the right of the heatmap), validating the identity of the populations. Gene ontology (GO) enrichment analysis of the differentially expressed genes revealed a distinct functional gene expression program for each macrophage population (**Fig. 3; C and D, table S2**). As expected, iNOS Mϕ are enriched for a pro-inflammatory program (*e.g.*, chemotaxis, positive regulation of the inflammatory process, including Nos2). iNOS Mϕ were also enriched for endocytosis and pinocytosis, including factors that regulate phagocytosis (e.g., Gas7 and Msr1, Galectins) and receptors that recognize microbial ligands (*e.g*., Dhx29, Clec4n, Siglec-1), indicating that these cells can readily ingest and potentially eliminate invading pathogens. In contrast, CD9 Mϕ showed enrichment for anti-inflammatory processes including cell proliferation, migration and negative regulation of the inflammatory process. Noteworthy, these processes included fatty acid and lipid metabolism (*e.g. Dgat2*, *Rara*, and *Sema4d*), beta-adrenergic signaling (*e.g*., *Adrb2*, *Adrbk2*) and prostaglandins degradation (*Hpgd)*, which can suggest the metabolic and activation states of these cells (Ağaç et al., 2018; Batista-Gonzalez et al., 2020). Importantly, these distinct activation programs indicated that CD9 Mϕ and iNOS Mϕ may have different functional roles during intracellular *S*.Tm infection.

**Figure 3:**
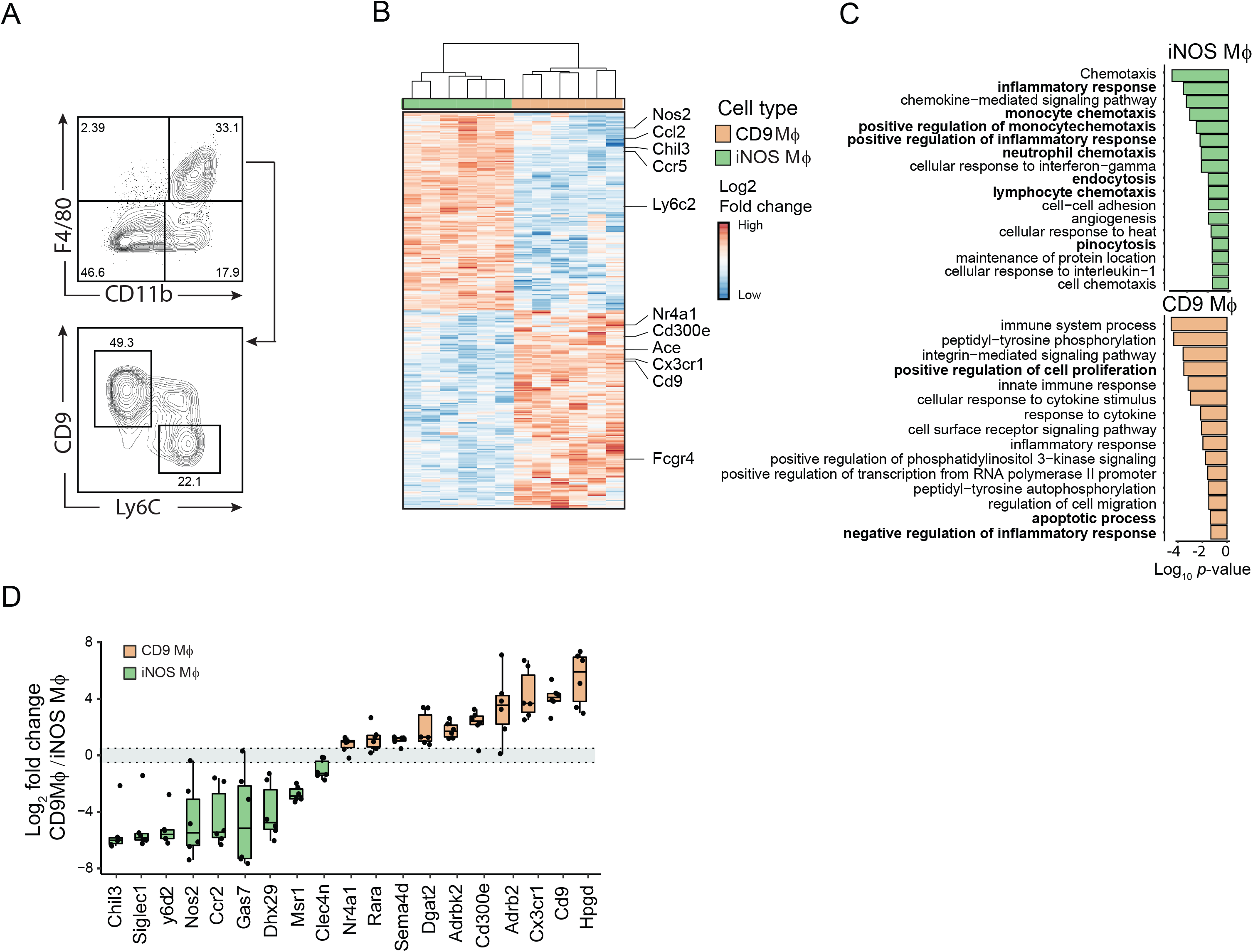
Functionally distinct activation programs of iNOS Mϕ and CD9 Mϕ. **(A)** Mice were infected with *S.*Tm and 24 hpi spleens were harvested, and analyzed by flow cytometry using CD11b, F4/80, CD9 and Ly6C antibodies. Presented are contour plots of iNOS Mϕ and CD9 Mϕ from the gated Cd11b^+^F4/80^+^ population. Results are representative of four mice. (**B-D**) Mice were infected as in A (n=6), and CD9 Mϕ and iNOS Mϕ were sorted by FACS and analyzed by bulk RNA-seq. Presented is a heatmap of the significant differentially expressed genes between CD9 Mϕ (orange) and iNOS Mϕ (green) (**B**), gene ontology annotations enriched in each population (**C**), and log2 fold expression changes of selected genes (grey area denotes borders of significant differences in expression, mice indicated by dots) (**D**).

### ELID was mirrored by a decline in Rp Mϕ, and increase in CD9 Mϕ

We then characterized the dynamics of the macrophage populations using flow cytometry, as well as their intracellular bacteria load using *S*.Tm-GFP (**Fig. 4; A and B, fig. S4; A-B**). In naïve mice, ~90% of the F4/80^+^ cells were Rp Mϕ. As infection progressed, Rp Mϕ abundance decreased and reached ~10% at 24 hpi. Conversely, iNOS Mϕ and CD9 Mϕ were almost undetectable in naïve mice and accumulated during the first hours until they reached at 24 hpi up to ~30% and ~55% of the splenic macrophage pool, respectively. Analysis of infected cells using the GFP signal of the intracellular bacteria revealed that during the first 8 hours *S*.Tm were found mostly within Rp Mϕ. In contrast, by 24 hpi, most *S*.Tm resided within CD9 Mϕ and iNOS Mϕ. The numeric increase of iNOS Mϕ and CD9 Mϕ coincided with intracellular growth of *S*.Tm during the second phase of the eclipse. To directly examine intracellular bacterial replication, we infected mice with *S*.Tm-GFP, and 24 hpi analyzed scCFU within CD9 Mϕ and iNOS Mϕ (**Fig. 4C**). In iNOS Mϕ, we measured no more than a single bacterium per cell. In contrast, 30% of CD9 Mϕ contained more than a single bacteria, suggesting that these cells are permissive to *S*.Tm intracellular growth. Using microscopy we confirmed that at 24 hpi 70% of the F4/80^+^CD9^+^ (CD9 Mϕ) contained more than three intracellular bacteria suggesting intracellular replication, compared to only 10% of the F4/80^+^CD9^−^ (**Fig. 4D**). This higher number of intracellular bacteria in CD9 Mϕ as evident by microscopy compared to scCFU may underscore a deficiency of *S*.Tm to establish re-growth following intracellular infection (Helaine et al., 2014).

**Figure 4:**
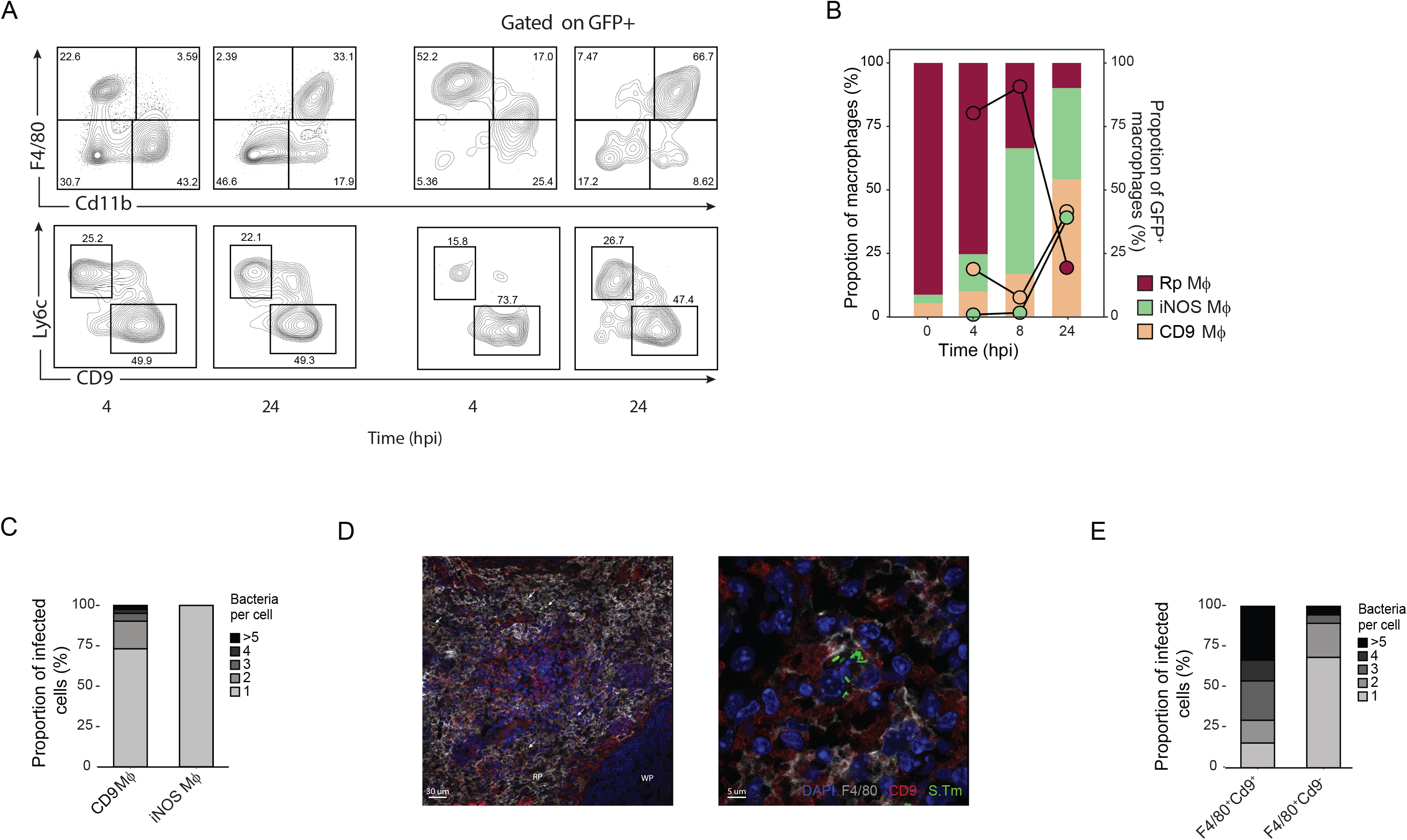
ELID was mirrored by a decline of Rp Mϕ and increase of CD9 Mϕ. (**A+B**) Mice were infected with *S.*Tm-GFP (n=3 per time point). At the indicated time points, spleens were harvested and analyzed by flow cytometry using antibodies to the indicated populations or the GFP signal of the bacteria (**A**). Bar graphs (left Y-axis) indicates mean ratio from the population of F4/80, and dots (right Y-axis) indicates cells containing intracellular bacteria (presented as ratio from F4/80^+^ GFP^+^ population) (**B**). (**C**) Mice were infected as in A and 24 hpi single GFP^+^ CD9 Mϕ and iNOS Mϕ were sorted and scCFU was measured (n=494, n=553 respectively). Presented are the proportions cells according to number of bacteria per cell (scCFU). (**D+E**) Mice were infected as in A. Twenty-four hpi spleens were harvested, fixed, stained and imaged with CD9 and F4/80 antibodies, DAPI and GFP for intracellular bacteria (**D**). Left image; gross morphology of the spleen with red pulp (RP) and white pulp (WP) indicated. Arrows indicate multiple intracellular *S*.Tm within CD9 Mϕ (F4/80^+^Cd9^+^). Right image; An example of CD9 Mϕ containing multiple bacteria. Number of intracellular bacteria were quantified across F4/80^+^Cd9^+^ or F4/80^+^Cd9^−^ cells (n=94 cells from 3 mice) (**E**).

Our results indicate that changes in the landscape of macrophage populations mirror the growth dynamics of *S*.Tm, and support a model where in the first phase of ELID, Rp Mϕ control and eliminate intracellular *S*.Tm, resulting in Rp Mϕ decline. Bacterial growth in the second expansion phase of ELID coincides with intracellular replication of *S*.Tm within the newly arising CD9 Mϕ.

### Rp Mϕ undergo necroptotic cell death and restricted bacterial growth during the first phase of the eclipse

Our observation that the majority of *S*.Tm were found within Rp Mϕ during the first phase of ELID led us to hypothesize that these cells mediate intracellular pathogen clearance, as was previously proposed (Borges da Silva et al., 2015). To test this, we utilized mice that harbor a diphtheria toxin receptor (DTR) under the promoter of CD169 that is expressed by Rp Mϕ (Miyake et al., 2007). CD169 ^DTR^ mice were treated with a single dose of either PBS (vehicle) or diphtheria toxin (DTx) to deplete Rp Mϕ, which was confirmed by flow cytometry analysis (*P*<0.05, **Fig. 5; A and B**). DTx treatment also significantly reduced the abundance of iNOS Mϕ, probably due to their low, but detectable expression of CD169 (**fig. S5A**). Next, CD169 ^DTR^ mice were treated with vehicle or DTx and infected with *S*.Tm for measurement of bacterial CFU (**Fig. 5C**). ELID of *S*.Tm was measured in vehicle-treated mice, but no CFU decline was evident in the DTx-treated animals in the first ELID phase (<8 hours). Kinetics of the second ELID phase (>8 hours) were similar between vehicle and DTx treated mice. Collectively, these data suggest that Rp Mϕ eliminate intracellular bacteria during the first phase of ELID.

**Figure 5:**
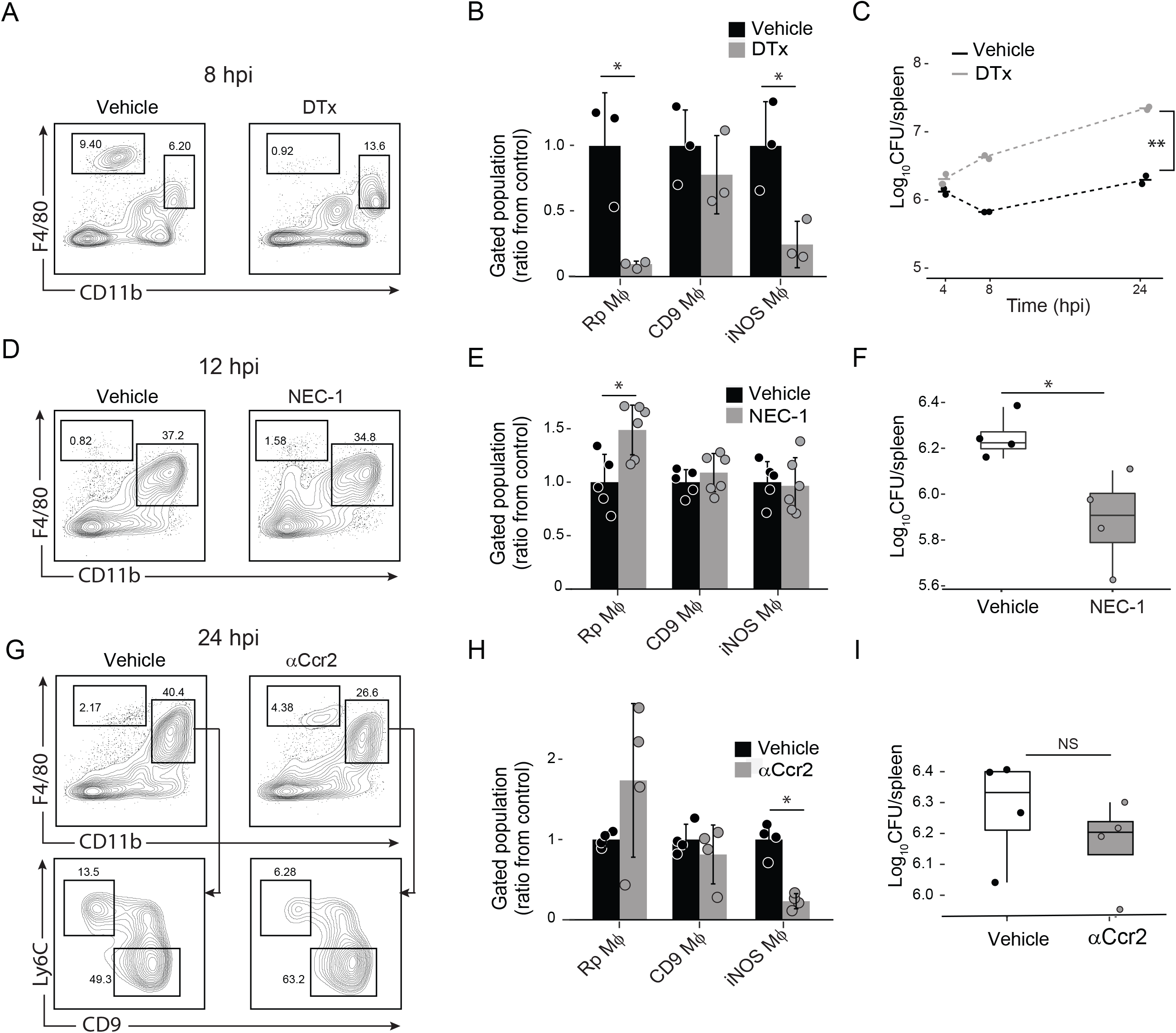
Rp Mϕ restricted *S.*Tm growth during the first phase of ELID. (**A+B**) Mice expressing diphtheria toxin receptor under siglec-1 promoter were treated with diphtheria toxin (DTx; n=3) or PBS (vehicle; n=3). Twenty-four hours later, mice were infected with *S.*Tm, and 8 hpi spleens were harvested. Spleens were analyzed by flow cytometry using antibodies to mark the indicated populations (**A**), and presented as the ratio of the population from vehicle-treated mice (**B**). (**C**) Mice were treated as in A, spleens were harvested at the indicated time points and CFU measured (n=2 per treatment in each time point). (**D-F**) Mice were injected i.v with either NEC-1 (n=5) or vehicle (n=4). One hour later, mice were infected with *S.*Tm, and 12 hpi, spleens were harvested. Spleens were analyzed by flow cytometry using antibodies to mark the indicated populations (**D**), presented as the ratio of the population from vehicle-treated mice (**E**) and plated for CFU (n=4 per treatment) (**F**). (**G-I**) Mice were injected intraperitoneally with αCcr2 antibodies or vehicle (n=4 per treatment). Twenty-four hours later, mice were infected with *S.*Tm, and 24 hpi spleens were harvested. Spleens were analyzed by flow cytometry using antibodies to mark the indicated populations (**G**), presented as the ratio of the population from vehicle-treated mice (**H**), and plated for CFU (**I**). [(**B**), (**E**) and (**H**)] Error bars indicate standard deviation. (**B**) * *P*<0.05, using student T-test. [(**E**), (**F**) and (**H**)] * *P*<0.05, using Mann–Whitney U test. (**C**) ** *P*<0.01 using two-way analysis of variance.

*Listeria monocytogenes* infection results in cell death of resident Kupfer cells in the liver through necroptosis (Blériot et al., 2015). We hypothesized that this may also be the mechanism of Rp Mϕ decline we observe in the first ELID phase (**Fig. 4A**). We therefore treated mice with the necroptosis inhibitor Necrostatin-1 (NEC-1) one hour before infection with *S*.Tm. At 12 hpi we measured significantly higher levels of Rp Mϕ in NEC-1 treated mice compared to vehicle-treated mice (**Fig. 5; D and E**). At 24 hpi we measured significantly lower bacterial load in NEC-1 treated mice compared to vehicle-treated mice (*P* < 0.05, **Fig. 5F**). These results suggest that *S*.Tm infection in the first phase of ELID is controlled by Rp Mϕ that then undergo necroptotic cell death.

During gut infection, inflammatory monocytes promote *S*.Tm expansion in the lumen of the inflamed intestine (McLaughlin et al., 2019). We turned to test the involvement of these cells in the second ELID phase. Classical monocyte migration from the bone marrow to infected tissues is dependent on Ccr2 (Boring et al., 1997), whereupon they differentiate to iNOS-producing cells and contribute to infection control (Tam et al., 2014). We specifically depleted classical monocytes by intraperitoneal injection of αCcr2 antibody (Brühl et al., 2007). Flow cytometry analysis of the spleen confirmed that αCcr2 treatment resulted in significantly reduced numbers of classical monocytes, as well as iNOS Mϕ, corroborating the precursor product relationship of these cells (*P*<0.05, **Fig. 5; G and H, fig. S5; B and C**). CD9 Mϕ and Rp Mϕ numbers remained unchanged. Moreover, treatment with αCcr2 did not significantly change bacterial CFU (**Fig. 5I**), suggesting that iNOS Mϕ have a limited contribution to *S*.Tm control during the second ELID phase.

### CD9 Mϕ originated from NC monocytes and provided an intracellular replication niche for *S*.Tm during the second phase of the eclipse

The above data suggest a critical role of CD9 Mϕ in *S*.Tm expansion. *k*NN-classification had indicated that CD9 Mϕ arise probably from NC monocytes (**Fig. 2E**), suggesting a possible approach for in vivo manipulation of this population. Specifically, ablation of NC monocytes can be achieved by a deletion of a super enhancer domain that controls NC expression of the survival factor Nr4a1, termed E2, without affecting Nr4a1 expression in other tissue macrophages (Thomas et al., 2016). Analysis of WT and *Nr4a1^e2−/−^* mice validated a significant decrease of NC monocytes in naïve mice (*P*<0.05, **Fig. 6; A and B**). We next infected WT and *Nr4a1^e2−/−^* mice with *S*.Tm. Compared to controls, *Nr4a1*^e2−/−^ mice displayed a significant decrease of CD9 Mϕ (*P*<0.05, **Fig. 6; C and D**). This provides an experimental confirmation that CD9 Mϕ are indeed macrophages derived from NC monocytes. In contrast and in line with their derivation from classical Ly6C^high^ monocytes, iNOS Mϕ were unaffected. Analysis of the infection kinetics and CFU measurements revealed that *Nr4a1^e2−/−^* mice displayed a significantly reduced splenic bacterial load, compared to WT mice (**Fig. 6E**). Finally, we infected WT or *Nr4a1^e2−/−^* mice with a low dose of *S*.Tm (1000 CFU) to test extended infection and survival of mice. Mutant mice lacking NC and CD9 Mϕ lost significantly less weight than controls (*P*<0.01, **Fig. 6F**) and displayed significantly extended survival compared to WT mice (*P*<0.01, **Fig. 6G**). Collectively, the lower CFU and extended survival in *Nr4a1^e2−/−^* mice, indicated that CD9 Mϕ are an intracellular niche required for *S*.Tm expansion, which has implication for the survival of the entire organism.

**Figure 6:**
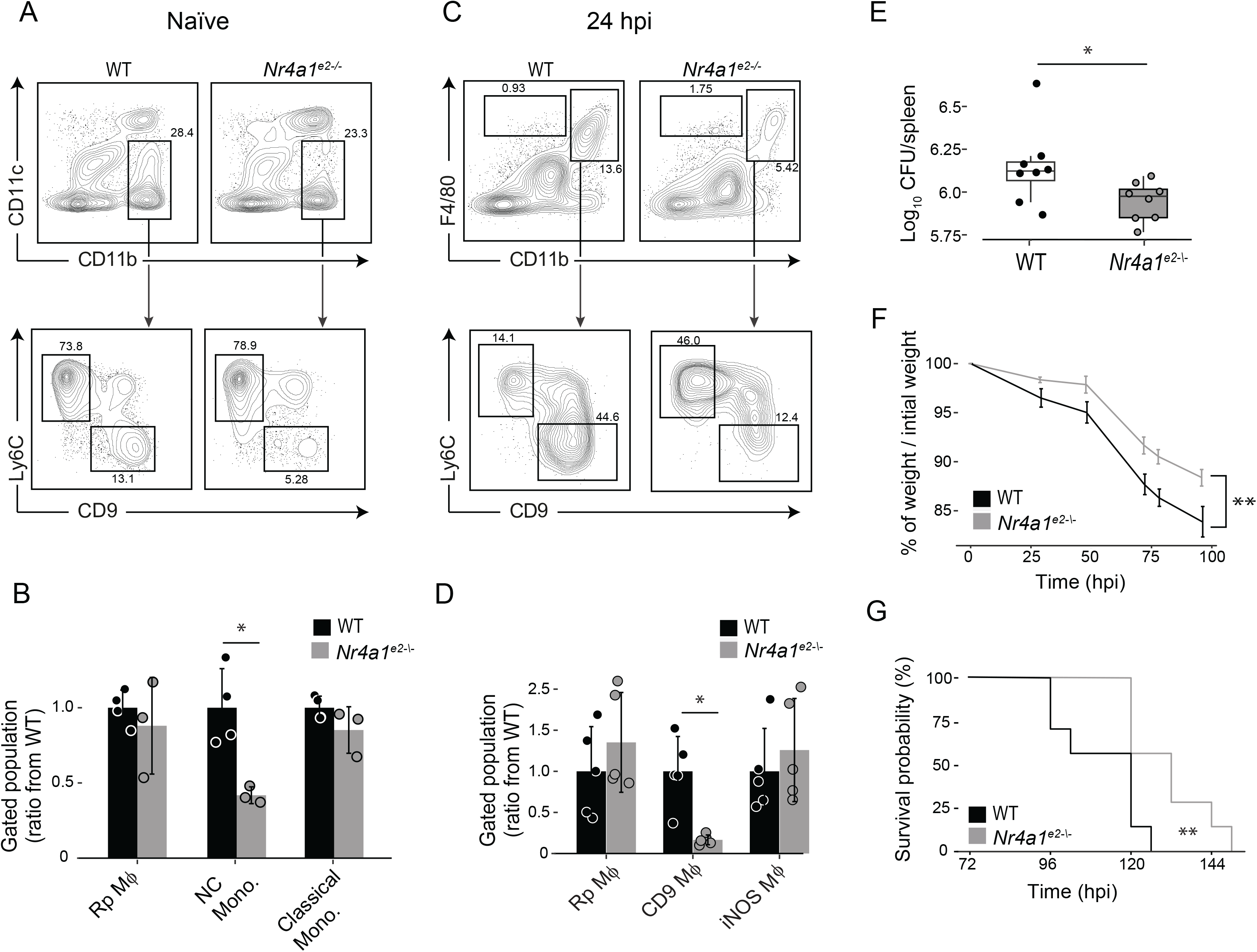
CD9 Mϕ provided a niche for intracellular *S*.Tm replication during the second phase of ELID. **(A+B)** Spleens of *Nr4a1^e2+/+^* (WT; n=3) and *Nr4a1^e2−/−^* (n=4) were harvested and analyzed by flow cytometry using antibodies to mark the indicated populations (**A**), and presented as the ratio of the population from WT mice (**B**). **(C-E)** WT (n=8) and *Nr4a1^e2−/−^* mice (n=8) were infected with *S.*Tm. Twenty-four hpi spleens were harvested and analyzed by flow cytometry using antibodies to mark the indicated populations (**C**), presented as the ratio of the population from WT mice (**D**) and plated for CFU (**E**). **(F+G)** WT (n=7) and *Nr4a1^e2−/−^* mice (n=7) were infected with 1000 CFU of *S.*Tm. Mice were weighed (**F**) and monitored for survival, presented on a Kaplan-Mayer plot (**G**). [(**B**) and (**D**)] Error bars indicated standard deviation. (**F**) Error bars indicated standard error. (**B**) * *P*<0.05 student t-test. [(**D**)+(**E**)] * *P*<0.05, Mann–Whitney U test. (**F**) ** *P*<0.01 two-way analysis of variance. (**G**) \*\**P*<0.01 log-rank test.

## Discussion

Mononuclear phagocytes can derive from several sources, including tissue-resident populations and infiltrating monocytes that differentiate into effector cells. At sites of infection, macrophages engage invading pathogens and induce effector processes to kill the invading agents. Intracellular bacterial pathogens present a paradox, as they preferably replicate within macrophages, the very cells that aim to destroy them (Price and Vance, 2014). Mostly, this duality was explained by differential signaling and gene expression programs of macrophages towards an inflammatory state (M1 that eliminates the pathogen) or an anti-inflammatory state (M2 that is more permissive for intracellular replication) (Pham et al., 2020; Saliba et al., 2017). Here, using scRNA-seq, we identified co-existing macrophage populations during early infection, each with unique molecular activation features, that concluded distinct outcomes of intracellular *S*.Tm infection. According to our results, during in vivo infection activation phenotypes of macrophages arise not from bifurcation of a macrophage population, but is rather a “division of labor” between different macrophage progeny that determines their activation state, as we show here for iNOS Mϕ from classical monocytes (M1-like) and CD9 Mϕ from NC monocytes (M2-like).

During early systemic infection, ELID was mediated by dispersal of *S*.Tm within three functionally distinct populations of F4/80 macrophages: tissue-resident Rp Mϕ, classical monocytes that give rise to iNOS Mϕ and CD9 Mϕ originated from NC monocytes. In the first phase of ELID, *S*.Tm is controlled by Rp Mϕ, at the expense of their demise. Most tissue resident macrophages are known to have limited microbicidal activity (Davies et al., 2013). It has been suggested that rather than directly eliminating invading bacteria, tissue resident macrophages undergo necroptotic cell death that triggers chemokine and cytokine release and recruitment of monocytes which eliminate the bacteria (Ginhoux et al., 2017). Indeed, inhibition of Rp Mϕ necroptotic cell death resulted in lower bacterial survival (**Fig. 5F**), which is in accordance with susceptibility to *S*.Tm infection of Rip3^−/−^ mice, that cannot undergo necroptotic cell death (Robinson et al., 2012).

In the second phase of ELID, intracellular *S*.Tm resided within iNOS Mϕ and CD9 Mϕ. A direct comparison between iNOS Mϕ and CD9 Mϕ indicated disparate molecular details and distinct phenotypic outcome of intracellular infection. Our understanding of the contribution of NC monocytes to tissue macrophage populations is limited. It was suggested that NC monocytes differentiate into alternatively activated macrophages (Narasimhan et al., 2019). Importantly, *S*.Tm growth during the second phase of the eclipse is mediated by replication within CD9 Mϕ (**Fig. 4**) and mice deficient of CD9 Mϕ alter the course of infection of the entire organism (**Fig. 6G**). Thus, a plausible model of ELID is that CD9 Mϕ engulf necroptotic Rp Mϕ which contain intracellular *S.*Tm. Thereby, *S*.Tm is delivered as a cargo to CD9 Mϕ, whereupon they can establish an intracellular niche (**fig. S6**).

We foresee that CD9 Mϕ may be relevant to other models of infection with *S*.Tm, including chronic infection and oral route administration. During chronic infection, *S.*Tm resides within macrophages recruited to granulomas (Goldberg et al., 2018), that are actively directed by *S*.Tm towards an anti-inflammatory phenotype (Pham et al., 2020). It will be interesting to test whether NC monocytes that give rise to CD9 Mϕ are involved in the anti-inflammatory phenotype of macrophages during chronic infection. During oral infection, *S*.Tm can traverse the epithelial barrier of the gut to the bloodstream and disseminate to the spleen either through invasion of epithelial cells (Frost et al., 1997; Takeuchi, 1967) or transported via CD18-expressing phagocytes (Vazquez-Torres et al., 1999). NC monocytes are also found within the gut (Schleier et al., 2020), and their contribution to macrophages in the gut mucosa is yet to be determined.

In summary, our work described that at early systemic infection, intracellular *S*.Tm is distributed within three distinct macrophage populations in the spleen, with diverse progeny and activation programs, that determines an eclipse growth dynamics. Beyond *S*.Tm infection, our model of a cell-type specific host-pathogen interaction may be a more general theme, with implications to infection with other intracellular pathogens.

## References

Ağaç, D., Estrada, L.D., Maples, R., Hooper, L.V., and Farrar, J.D. (2018). The β2-adrenergic receptor controls inflammation by driving rapid IL-10 secretion. Brain. Behav. Immun. 74, 176–185.

Baran, Y., Bercovich, A., Sebe-Pedros, A., Lubling, Y., Giladi, A., Chomsky, E., Meir, Z., Hoichman, M., Lifshitz, A., and Tanay, A. (2019). MetaCell: analysis of single-cell RNA-seq data using K-nn graph partitions. Genome Biol. 20, 206.

Batista-Gonzalez, A., Vidal, R., Criollo, A., and Carreño, L.J. (2020). New Insights on the Role of Lipid Metabolism in the Metabolic Reprogramming of Macrophages. Front. Immunol. 10.

Blériot, C., Dupuis, T., Jouvion, G., Eberl, G., Disson, O., and Lecuit, M. (2015). Liver-Resident Macrophage Necroptosis Orchestrates Type 1 Microbicidal Inflammation and Type-2-Mediated Tissue Repair during Bacterial Infection. Immunity 42, 145–158.

Borges da Silva, H., Fonseca, R., Pereira, R.M., Cassado, A. dos A., Álvarez, J.M., and D’Império Lima, M.R. (2015). Splenic Macrophage Subsets and Their Function during Blood-Borne Infections. Front. Immunol. 6.

Boring, L., Gosling, J., Chensue, S.W., Kunkel, S.L., Farese, R.V., Broxmeyer, H.E., and Charo, I.F. (1997). Impaired monocyte migration and reduced type 1 (Th1) cytokine responses in C-C chemokine receptor 2 knockout mice. J. Clin. Invest. 100, 2552–2561.

Brühl, H., Cihak, J., Plachý, J., Kunz-Schughart, L., Niedermeier, M., Denzel, A., Gomez, M.R., Talke, Y., Luckow, B., Stangassinger, M., et al. (2007). Targeting of Gr-1+,CCR2+ monocytes in collagen-induced arthritis. Arthritis Rheum. 56, 2975–2985.

Carter, P.B., and Collins, F.M. (1974). The route of enteric infection in normal mice. J. Exp. Med. 139, 1189–203.

Chakarov, S., Lim, H.Y., Tan, L., Lim, S.Y., See, P., Lum, J., Zhang, X.-M., Foo, S., Nakamizo, S., Duan, K., et al. (2019). Two distinct interstitial macrophage populations coexist across tissues in specific subtissular niches. Science 363.

Cirillo, D.M., Valdivia, R.H., Monack, D.M., and Falkow, S. (1998). Macrophage-dependent induction of the Salmonella pathogenicity island 2 type III secretion system and its role in intracellular survival. Mol. Microbiol. 30, 175–188.

Davies, L.C., Jenkins, S.J., Allen, J.E., and Taylor, P.R. (2013). Tissue-resident macrophages. Nat. Immunol. 14, 986–995.

De Jesus, M., Park, C.G., Su, Y., Goldman, D.L., Steinman, R.M., and Casadevall, A. (2008). Spleen deposition of Cryptococcus neoformans capsular glucuronoxylomannan in rodents occurs in red pulp macrophages and not marginal zone macrophages expressing the C-type lectin SIGN-R1. Med. Mycol. 46, 153–162.

Dick, S.A., Macklin, J.A., Nejat, S., Momen, A., Clemente-Casares, X., Althagafi, M.G., Chen, J., Kantores, C., Hosseinzadeh, S., Aronoff, L., et al. (2019). Self-renewing resident cardiac macrophages limit adverse remodeling following myocardial infarction. Nat. Immunol. 20, 29.

Ercoli, G., Fernandes, V.E., Chung, W.Y., Wanford, J.J., Thomson, S., Bayliss, C.D., Straatman, K., Crocker, P.R., Dennison, A., Martinez-Pomares, L., et al. (2018). Intracellular replication of Streptococcus pneumoniae inside splenic macrophages serves as a reservoir for septicaemia. Nat. Microbiol. 3, 600–610.

Frost, A.J., Bland, A.P., and Wallis, T.S. (1997). The early dynamic response of the calf ileal epithelium to Salmonella typhimurium. Vet. Pathol. 34, 369–386.

Geddes, K., Iii, F.C., and Heffron, F. (2007). Analysis of Cells Targeted by Salmonella Type III Secretion In Vivo. PLOS Pathog. 3, e196.

Giladi, A., Wagner, L.K., Li, H., Dörr, D., Medaglia, C., Paul, F., Shemer, A., Jung, S., Yona, S., Mack, M., et al. (2020). Cxcl10 + monocytes define a pathogenic subset in the central nervous system during autoimmune neuroinflammation. Nat. Immunol. 21, 525–534.

Ginhoux, F., and Guilliams, M. (2016). Tissue-Resident Macrophage Ontogeny and Homeostasis. Immunity 44, 439–449.

Ginhoux, F., Bleriot, C., and Lecuit, M. (2017). Dying for a Cause: Regulated Necrosis of Tissue-Resident Macrophages upon Infection. Trends Immunol. 38, 693–695.

Goldberg, M.F., Roeske, E.K., Ward, L.N., Pengo, T., Dileepan, T., Kotov, D.I., and Jenkins, M.K. (2018). Salmonella Persist in Activated Macrophages in T Cell-Sparse Granulomas but Are Contained by Surrounding CXCR3 Ligand-Positioned Th1 Cells. Immunity.

Guilliams, M., Mildner, A., and Yona, S. (2018). Developmental and Functional Heterogeneity of Monocytes. Immunity 49, 595–613.

Hashimshony, T., Wagner, F., Sher, N., and Yanai, I. (2012). CEL-Seq: single-cell RNA-Seq by multiplexed linear amplification. Cell Rep 2, 666–673.

Helaine, S., Cheverton, A.M., Watson, K.G., Faure, L.M., Matthews, S.A., and Holden, D.W. (2014). Internalization of Salmonella by macrophages induces formation of nonreplicating persisters. Science 343, 204–208.

Hensel, M., Shea, J.E., Waterman, S.R., Mundy, R., Nikolaus, T., Banks, G., Vazquez-Torres, A., Gleeson, C., Fang, F.C., and Holden, D.W. (1998). Genes encoding putative effector proteins of the type III secretion system of Salmonella pathogenicity island 2 are required for bacterial virulence and proliferation in macrophages. Mol. Microbiol. 30, 163–174.

Huang, L., Nazarova, E.V., Tan, S., Liu, Y., and Russell, D.G. (2018). Growth of Mycobacterium tuberculosis in vivo segregates with host macrophage metabolism and ontogeny. J. Exp. Med. jem.20172020.

Italiani, P., and Boraschi, D. (2014). From Monocytes to M1/M2 Macrophages: Phenotypical vs. Functional Differentiation. Front. Immunol. 5.

Jaitin, D.A., Kenigsberg, E., Keren-Shaul, H., Elefant, N., Paul, F., Zaretsky, I., Mildner, A., Cohen, N., Jung, S., Tanay, A., et al. (2014). Massively parallel single cell RNA-Seq for marker-free decomposition of tissues into cell types. Science 343, 776–779.

Jaitin, D.A., Adlung, L., Thaiss, C.A., Weiner, A., Li, B., Descamps, H., Lundgren, P., Bleriot, C., Liu, Z., Deczkowska, A., et al. (2019). Lipid-Associated Macrophages Control Metabolic Homeostasis in a Trem2-Dependent Manner. Cell 178, 686–698.e14.

Kirby, A.C., Beattie, L., Maroof, A., van Rooijen, N., and Kaye, P.M. (2009). SIGNR1-Negative Red Pulp Macrophages Protect against Acute Streptococcal Sepsis after Leishmania donovani-Induced Loss of Marginal Zone Macrophages. Am. J. Pathol. 175, 1107–1115.

Kolter, J., Feuerstein, R., Zeis, P., Hagemeyer, N., Paterson, N., d’Errico, P., Baasch, S., Amann, L., Masuda, T., Lösslein, A., et al. (2019). A Subset of Skin Macrophages Contributes to the Surveillance and Regeneration of Local Nerves. Immunity 50, 1482–1497.e7.

McLaughlin, P.A., Bettke, J.A., Tam, J.W., Leeds, J., Bliska, J.B., Butler, B.P., and van der Velden, A.W.M. (2019). Inflammatory monocytes provide a niche for Salmonella expansion in the lumen of the inflamed intestine. PLOS Pathog. 15, e1007847.

Menezes, S., Melandri, D., Anselmi, G., Perchet, T., Loschko, J., Dubrot, J., Patel, R., Gautier, E.L., Hugues, S., Longhi, M.P., et al. (2016). The Heterogeneity of Ly6Chi Monocytes Controls Their Differentiation into iNOS+ Macrophages or Monocyte-Derived Dendritic Cells. Immunity 45, 1205–1218.

Mills, E., and Avraham, R. (2017). Breaking the population barrier by single cell analysis: one host against one pathogen. Curr. Opin. Microbiol. 36, 69–75.

Miyake, Y., Asano, K., Kaise, H., Uemura, M., Nakayama, M., and Tanaka, M. (2007). Critical role of macrophages in the marginal zone in the suppression of immune responses to apoptotic cell–associated antigens. J. Clin. Invest. 117, 2268–2278.

Narasimhan, P.B., Marcovecchio, P., Hamers, A.A.J., and Hedrick, C.C. (2019). Nonclassical Monocytes in Health and Disease. Annu. Rev. Immunol. 37, 439–456.

Olingy, C.E., San Emeterio, C.L., Ogle, M.E., Krieger, J.R., Bruce, A.C., Pfau, D.D., Jordan, B.T., Peirce, S.M., and Botchwey, E.A. (2017). Non-classical monocytes are biased progenitors of wound healing macrophages during soft tissue injury. Sci. Rep. 7, 447.

Pham, T.H.M., Brewer, S.M., Thurston, T., Massis, L.M., Honeycutt, J., Lugo, K., Jacobson, A.R., Vilches-Moure, J.G., Hamblin, M., Helaine, S., et al. (2020). Salmonella-Driven Polarization of Granuloma Macrophages Antagonizes TNF-Mediated Pathogen Restriction during Persistent Infection. Cell Host Microbe 27, 54–67.e5.

Price, J.V., and Vance, R.E. (2014). The Macrophage Paradox. Immunity 41, 685–693.

Robinson, N., McComb, S., Mulligan, R., Dudani, R., Krishnan, L., and Sad, S. (2012). Type I interferon induces necroptosis in macrophages during infection with *Salmonella enterica* serovar Typhimurium. Nat. Immunol. 13, 954–962.

Salcedo, S.P., Noursadeghi, M., Cohen, J., and Holden, D.W. (2001). Intracellular replication of Salmonella typhimurium strains in specific subsets of splenic macrophages in vivo. Cell. Microbiol. 3, 587–597.

Saliba, A.-E., Li, L., Westermann, A.J., Appenzeller, S., Stapels, D.A.C., Schulte, L.N., Helaine, S., and Vogel, J. (2017). Single-cell RNA-seq ties macrophage polarization to growth rate of intracellular *Salmonella*. Nat. Microbiol. 2, 16206.

Schleier, L., Wiendl, M., Heidbreder, K., Binder, M.-T., Atreya, R., Rath, T., Becker, E., Schulz-Kuhnt, A., Stahl, A., Schulze, L.L., et al. (2020). Non-classical monocyte homing to the gut via α4β7 integrin mediates macrophage-dependent intestinal wound healing. Gut 69, 252–263.

Schyns, J., Bai, Q., Ruscitti, C., Radermecker, C., Schepper, S.D., Chakarov, S., Farnir, F., Pirottin, D., Ginhoux, F., Boeckxstaens, G., et al. (2019). Non-classical tissue monocytes and two functionally distinct populations of interstitial macrophages populate the mouse lung. Nat. Commun. 10, 1–16.

Serbina, N.V., Salazar-Mather, T.P., Biron, C.A., Kuziel, W.A., and Pamer, E.G. (2003). TNF/iNOS-producing dendritic cells mediate innate immune defense against bacterial infection. Immunity 19, 59–70.

Serbina, N.V., Jia, T., Hohl, T.M., and Pamer, E.G. (2008). Monocyte-Mediated Defense Against Microbial Pathogens. Annu. Rev. Immunol. 26, 421–452.

Swirski, F.K., Nahrendorf, M., Etzrodt, M., Wildgruber, M., Cortez-Retamozo, V., Panizzi, P., Figueiredo, J.-L., Kohler, R.H., Chudnovskiy, A., Waterman, P., et al. (2009). Identification of Splenic Reservoir Monocytes and Their Deployment to Inflammatory Sites. Science 325, 612–616.

Takeuchi, A. (1967). Electron microscope studies of experimental Salmonella infection. I. Penetration into the intestinal epithelium by Salmonella typhimurium. Am. J. Pathol. 50, 109–136.

Tam, J.W., Kullas, A.L., Mena, P., Bliska, J.B., and van der Velden, A.W.M. (2014). CD11b+ Ly6Chi Ly6G− Immature Myeloid Cells Recruited in Response to Salmonella enterica Serovar Typhimurium Infection Exhibit Protective and Immunosuppressive Properties. Infect. Immun. 82, 2606–2614.

Thomas, G.D., Hanna, R.N., Vasudevan, N.T., Hamers, A.A., Romanoski, C.E., McArdle, S., Ross, K.D., Blatchley, A., Yoakum, D., Hamilton, B.A., et al. (2016). Deleting an Nr4a1 Super-Enhancer Subdomain Ablates Ly6Clow Monocytes while Preserving Macrophage Gene Function. Immunity 45, 975–987.

Trzebanski, S., and Jung, S. (2020). Plasticity of monocyte development and monocyte fates. Immunol. Lett. 227, 66–78.

Vazquez-Torres, A., Jones-Carson, J., Bäumler, A.J., Falkow, S., Valdivia, R., Brown, W., Le, M., Berggren, R., Parks, W.T., and Fang, F.C. (1999). Extraintestinal dissemination of Salmonella by CD18-expressing phagocytes. Nature 401, 804–808.

Yona, S., Kim, K.-W., Wolf, Y., Mildner, A., Varol, D., Breker, M., Strauss-Ayali, D., Viukov, S., Guilliams, M., Misharin, A., et al. (2013). Fate Mapping Reveals Origins and Dynamics of Monocytes and Tissue Macrophages under Homeostasis. Immunity 38, 79–91.

